# GPR180 deficiency impairs mitochondrial function and insulin secretion in pancreatic β-cells

**DOI:** 10.64898/2026.04.23.720098

**Authors:** Matus Antal, Tina Dahlby, Peter Makovicky, Anastasiia Novak, Carla Horvath, Daniela Stanikova, Simona Gazova, Radana Brumarova, Eliska Ivanovova, Martina Horejsova, David Friedecky, Olga Krizanova, Marta Novotova, Daniela Gasperikova, Christian Wolfrum, Miroslav Balaz, Lucia Balazova

**Affiliations:** Institute of Experimental Endocrinology, Biomedical Research Center, Slovak Academy of Sciences, Bratislava, Slovakia; Institute of Food, Nutrition and Health, ETH Zurich, Schwerzenbach, Switzerland; Institute of Experimental Oncology, Biomedical Research Center, Slovak Academy of Sciences, Bratislava, Slovakia; Department of Pediatrics, Medical Faculty of Comenius University and National Institute for Children’s Diseases, Bratislava, Slovakia; Institute of Clinical and Translational Research, Biomedical Research Center, Slovak Academy of Sciences, Bratislava, Slovakia; Laboratory for Inherited Metabolic Disorders, Department of Clinical Biochemistry, University Hospital Olomouc and Faculty of Medicine and Dentistry, Palacky University Olomouc, Olomouc, Czechia; Department of Animal Physiology and Ethology, Faculty of Natural Sciences, Comenius University, Bratislava, Slovakia

**Keywords:** GPR180, pancreatic β-cells, insulin secretion, mitochondrial metabolism, β-cell identity

## Abstract

**Objective:** G protein-coupled receptor 180 (GPR180) has been implicated in systemic energy metabolism, primarily in adipose tissue and the liver. Given impaired whole-body glucose tolerance following GPR180 dysfunction, we aimed to determine whether GPR180 regulates pancreatic β-cell function. We investigated whether GPR180 contributes to β-cell insulin secretion by modulating metabolic processes that couple glucose sensing to mitochondrial energy production.

**Methods:** Phenotyping of whole-body (*Gpr180 ^-/-^*) and β cell-specific *Gpr180* (b*Gpr180*-KO) knockout mice was combined with gain- and loss-of-function studies in MIN6 cells. Glucose-stimulated insulin secretion, pancreatic endocrine architecture and identity, transcriptomic and metabolic profiles, as well as mitochondrial function were assessed using *in vivo* and *in vitro* approaches, including metabolic challenge tests, histology, RNA sequencing, targeted metabolomics, respirometry, and transmission electron microscopy.

**Results:** Loss of GPR180 impaired first-phase insulin secretion and glucose tolerance without affecting insulin sensitivity. These defects were β-cell-autonomous, as confirmed in the b*Gpr180*-KO mice and in MIN6 cells. Functional studies revealed that GPR180 regulates mitochondrial substrate utilization, anaplerotic support of the TCA cycle, and ATP generation without affecting glucose uptake or mitochondrial biogenesis. In particular, Gpr180-deficient β cells showed mitochondrial membrane depolarization, reduced oxygen consumption, and endoplasmic reticulum remodeling, altering the local mitochondrial microenvironment. *In vivo*, *Gpr180* deletion in β cells led to downregulation of mitochondrial gene programs in islets, along with altered endocrine cell identity.

**Conclusions:** GPR180 is a previously unrecognized regulator of pancreatic β-cell metabolic competence and identity, linking defects in insulin secretion with alterations in mitochondrial function and endocrine cell identity.

**Graphical abstract:** 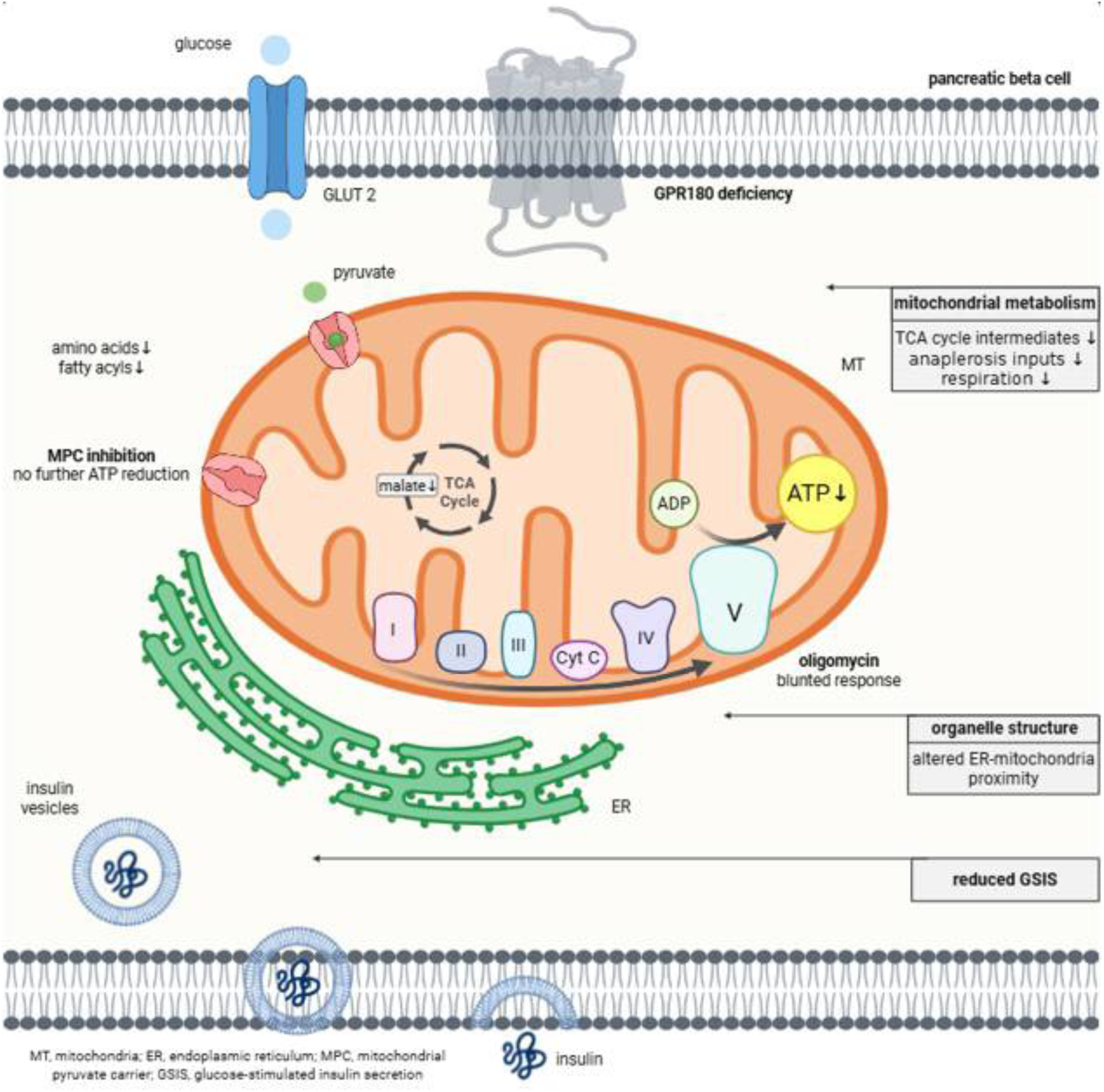

## Introduction

Pancreatic β-cells maintain glucose homeostasis by coupling nutrient metabolism to insulin secretion. Central to this process is mitochondrial oxidative metabolism, which links glucose-derived substrate flux to ATP production and subsequent insulin granule exocytosis [1]. Impairment of this metabolic coupling is a key contributor to β-cell dysfunction and the pathogenesis of diabetes [2]. Under metabolic stress, β-cells adapt by increasing insulin secretion and cellular mass to compensate for insulin resistance; however, this compensatory capacity is limited, leading to progressive β-cell failure [3]. Understanding the mechanisms that preserve β-cell functional identity and metabolic competence is therefore critical for maintaining glucose homeostasis [4].

Efficient insulin secretion requires tight coordination between glucose metabolism and mitochondrial function, but the upstream regulators that maintain this metabolic coupling remain incompletely defined [5]. While core mechanisms are established, how mitochondrial metabolism is precisely regulated to support β-cell function under stress or disease is still not fully resolved [4–6]. This underscores the need for identifying additional regulators of β-cell metabolic competence.

GPR180 is a seven-transmembrane domain protein with emerging roles in metabolic regulation. Previous studies have implicated GPR180 in the regulation of systemic energy balance, particularly in adipose tissue and the liver, as its loss affects glucose homeostasis and lipid metabolism [7–9]. Notably, global deletion of GPR180 leads to impaired glucose tolerance, whereas adipocyte-specific deletion only partially recapitulates this phenotype, suggesting contributions from additional tissues [7]. Despite these observations, the role of GPR180 in pancreatic β-cells and its potential involvement in β-cell metabolic regulation remain unknown. Given the central role of mitochondrial metabolism in β-cell function and the emerging role of GPR180 in systemic metabolic regulation, this study aimed to define GPR180’s role in β-cell function. Using complementary in vivo and in vitro approaches, we assessed the consequences of GPR180 deficiency on insulin secretion, mitochondrial metabolism, and endocrine cell identity.

## Methods

### Animal Models and Experimental Procedures

Global *Gpr180* knockout mice were generated via CRISPR/Cas9-mediated genome editing as described previously [7]. To obtain experimental cohorts, heterozygous mice were interbred to produce wild-type (WT) and knockout littermates. β-cell-specific *Gpr180* knockout (*Gpr180*^fl/fl^ x Rip-Cre, here b*Gpr180*-KO) mice were obtained by crossing *Gpr180*^fl/fl^ mice [7] with rat insulin promoter-driven Cre recombinase (RIP-Cre) transgenic mice kindly provided by Prof. Stoffel (ETH Zurich). Littermates lacking the RIP-Cre allele were used as controls. Experimental and control animals were kept mixed since weaning and treated/analysed in random order to minimise potential confounders.

All experiments were conducted on 12-14 week-old male mice. All mice had a C57BL/6N background. Animals were group-housed (3-4 mice) at 22°C on a 12-hour reversed light/dark cycle with *ad libitum* access to chow and water. To induce obesity, mice received a 60% fat diet (Provimi Kliba SA) from 14 weeks of age for 12 weeks. Animals were euthanized singly in a CO_2_ atmosphere. Animal experiments were approved by the Veterinary Office of the Canton of Zürich (Switzerland) and conducted in accordance with FELASA and institutional guidelines.

#### Glucose and Insulin Tolerance Test

After 6 hours of fasting, mice received i.p. injections of D-glucose (1 mg/g body weight) or insulin (0.75 U/kg). Tail-vein blood glucose was measured using a glucometer (ACCU-CHEK Aviva, Roche).

### Glucose-Stimulated Insulin Secretion

Mice were fasted for 6 hours and injected with glucose (1 mg/g i.p.). Blood samples (baseline and 5 minutes post-glucose) were collected in EDTA-coated tubes, centrifuged, and plasma insulin was measured by Ultrasensitive mouse insulin ELISA (Crystal Chem).

#### Isolation of Langerhans islets

Pancreatic islets were isolated via bile duct collagenase perfusion [10]. Tissue digestion (37°C/30 min) was followed by centrifugation at 567g/2 minutes/4°C, washing, and filtration sequentially through 500 μm and 70 μm nylon strainers. Overnight-cultured islets (RPMI-1640 with GlutaMax, 1% P/S, and 10% FBS; Gibco) were hand-picked and snap-frozen for analyses.

### Tissue processing

#### Insulin content

Insulin content was quantified from the whole pancreas using acid-ethanol extraction (1.5% HCl in 70% ethanol + 0.1% Triton X-100; 20 µL/mg tissue) followed by Ultrasensitive mouse insulin ELISA (Crystal Chem). Insulin content was normalized to pancreas weight.

#### Immunohistochemistry

The pancreas was fixed in 4% paraformaldehyde, washed in 70% ethanol, and embedded in paraffin blocks. Sections (5 µm) were obtained from the tail region of the pancreas and stained with haematoxylin-eosin or anti-Insulin antibody (1:1000, RRID: AB_3712125, Elabscience) followed by a secondary antibody conjugated to Alexa Fluor 488 (RRID: AB_143165, ThermoFisher Scientific). Imaging was performed using an Axiovert microscope (Zeiss). Islet morphometry was performed using Zen 3.9 software (Zeiss).

#### Gene expression analysis

Total RNA was extracted by phenol-chloroform, followed by DNase treatment. RNA quantity was measured by Qubit fluorometry (Invitrogen), and integrity was confirmed with an Agilent 2100 Bioanalyzer.

Knockout efficiency in isolated islets was confirmed by qPCR. First, extracted total RNA was reversed transcribed using the High Capacity cDNA Reverse transcription kit (Applied Biosystems), with 200 ng of RNA. Real-time PCR was performed using Fast SYBR Green Master Mix (Applied Biosystems) on a QuantStudio 5 system (ThermoFisher Scientific). Gene expression of *Gpr180* was normalized to housekeeping gene *Tbp*. All primer sequences are provided in Supplementary Table 1.

For mitochondrial DNA copy number, genomic DNA was extracted from the organic phase following RNA isolation from pancreatic islets using Back Extraction Buffer (4M GTC, 50 mM Na₃C₆H₅O₇, 1M Tris). DNA concentration and purity were measured spectrophotometrically with a Nanodrop (ThermoFisher Scientific), and samples were diluted to a uniform concentration prior to analysis. Relative mitochondrial DNA content was determined by normalizing expression of the mitochondrial gene *Nd1* to the nuclear reference gene *Lpl*.

Libraries for bulk RNA sequencing were prepared from >100 ng input RNA using poly(A) enrichment and directional library construction. mRNA sequencing was performed by Novogene Co., Ltd. (Munich, Germany) on an Illumina NovaSeq 6000 platform with paired-end 150 bp reads. Data output was ≥ 20 million read pairs per sample, ensuring comprehensive transcriptome coverage. Raw sequencing data were processed using Novogene’s in-house bioinformatics pipeline, NovoMagic, which includes quality filtering, alignment to the reference genome, transcript assembly, gene expression quantification, and differential expression analysis. Differentially expressed genes (DEGs) were defined by |log2(Fold Change)| ≥ 1 and adjusted p-value (padj) ≤ 0.05. Functional enrichment analyses (GO and KEGG) applied a significance threshold of padj < 0.05. We acknowledge Novogene for providing high-quality sequencing and bioinformatics services.

Gene expression analysis in MIN6 cells (standard culture conditions), particularly *t*otal RNA extraction, qPCR to determine knockdown as well as overexpression efficiency, mitochondrial copy number determination and RNAseq was performed as described above for isolated islets.

### Functional *in vitro* studies

#### Cell culture and Gpr180 modulation

MIN6 cells (passage 14-20, AddexBio) were cultured in 4.5g/l glucose DMEM-GlutaMAX medium with pyruvate (Gibco), 15% FBS (Gibco), 50 µM 2-mercaptoethanol, and 1% penicillin-streptomycin. For *Gpr180* silencing, cells were transfected with either 50 nM *Gpr180*-targeting siRNA (ID: 81632) or a non-targeting control siRNA (Silencer™ Select, Life Technologies) using Lipofectamine RNAiMAX (Invitrogen). Overexpression was performed by Turbofect (ThermoFisher Scientific) mediated transfection of the pcDNA3-smURFP-IRES-eGFP plasmid [11] (#80343, Addgene). The *GPR180* coding sequence was subcloned using XhoI and XbaI restriction sites. All downstream analyses were performed 48 hours post-transfection. Cell culture experiments were independently reproduced and were mycoplasma-free.

#### Glucose-stimulated insulin secretion

After 1-hour preincubation in low-glucose (2 mM) KRB buffer (129 mM NaCl, 4.8 mM KCl, 1 mM CaCl_2_, 5 mM NaHCO_3_, 1.2 mM KH_2_PO4, 1.2 mM MgSO_4_, 10 mM HEPES, 0.1% BSA, pH 7.4), cells were exposed to 2 mM or 25 mM glucose for 1 hour [12]. Secreted insulin was assessed (Ultrasensitive mouse insulin ELISA, Crystal Chem) and normalized to cellular proteins (DC assay). Cellular insulin content was measured following acid–ethanol extraction and quantified by ELISA.

#### Metabolomics

MIN6 cells cultured in standard medium 48h post-transfection were quenched and deproteinized on dry ice by adding 200 µL of ice-cold 80% methanol containing internal standards (10 µM creatinine-D3, 10 µM leucine-D3, 0.5 µM methylmalonate-D3, 0.1 µM butyrylcarnitine-D3, and 45 µM lactate-D3), followed by vortex mixing and sonication. After overnight protein precipitation at −80 °C, the samples were briefly vortexed and centrifuged. The supernatant from each sample was split into two aliquots: one part was used for wide- coverage targeted metabolomic analysis, and the other portion was evaporated to dryness and reconstituted in water for organic acid (OA) analysis. Equal aliquots from all samples were pooled to prepare the quality control (QC) sample. Wide-coverage targeted metabolomic analysis and organic acid analysis were performed using a Nexera LC system (Sciex, Framingham, MA, USA) coupled to a QTRAP 6500+ mass spectrometer (Sciex, Framingham, MA, USA). Both methods were carried out in positive and negative ion modes with polarity switching using scheduled multiple reaction monitoring (sMRM). Detailed information of the targeted metabolomic [13], and organic acid [14] methods, including chromatographic conditions and mass spectrometry settings, are provided in our previously published work.

Raw data from the metabolomic and OA analyses were acquired using SCIEX OS (v4.0) and Analyst software (v1.7.3) (SCIEX, Framingham, MA, USA). Data processing and statistical evaluation were performed in R (v4.5.1, www.r-project.org) using the Metabol package [15]. Local estimated smoothing signal (LOESS) correction was applied to each dataset independently. Metabolites with a coefficient of variation greater than 30% in QC samples were excluded from further analysis. The datasets were merged prior to statistical evaluation, and a natural logarithmic transformation was applied. Results were visualized using box plots, a Cytoscape metabolic map (Cytoscape v3.10.4, Bethesda, MD, USA), and MetaboAnalyst software (v6.0) [16]. Within Cytoscape, node size corresponded to the −log(p-value) derived from the Student’s t-test, whereas node color indicated the fold change, with red denoting an increase and blue a decrease in GPR180-deficient cells relative to controls. In MetaboAnalyst, Pareto scaling was applied to the data structure. In the pathway analysis, the x-axis represents the pathway impact score, whereas the y-axis shows the −log(p-value) obtained from the enrichment analysis. The enrichment analysis generated a ranked set of pathways affected in the dataset, visualizing the top 25 pathways according to their statistical relevance.

#### Glucose Uptake and Glucokinase Activity

Glucose uptake was quantified using the Glucose Uptake-Glo™ Assay Kit (Promega). Glucokinase activity was assessed by the Glucokinase Activity Fluorometric Assay Kit (Abcam). Luminescence and fluorescence were measured using a SynergyMx Plate reader, and the data were analysed by Gen5 v3.08 software (BioTek).

#### Mitochondrial phenotyping

Mitochondrial mass was evaluated using two approaches: the *Nd1/Lpl* ratio as described above and staining of mitochondria using MitoTracker™ Green (ThermoFisher Scientific).

Mitochondrial membrane potential was assessed using the JC-1 Assay Kit (ThermoFisher Scientific).

Mitochondrial respiration was assessed by the Seahorse XF Pro Analyzer (Agilent). MIN6 cells were plated, and transfections were performed on 96-well Seahorse microplates. On the day of the experiment, the standard culture medium was replaced with XF DMEM Medium (pH 7.4, Agilent) supplemented with glucose (25 mM; Sigma–Aldrich), 2 mM sodium pyruvate (Invitrogen), and 2 mM GlutaMax (Invitrogen). Test compounds were sequentially injected to achieve the following final concentrations: 1.25 µM Oligomycin, 10 µM FCCP, 3 µM Rotenone with 3.75 µM Antimycin A. All compounds were purchased from Sigma–Aldrich. The oxygen consumption rate (OCR) was normalized to protein content per well (µg protein). Non-mitochondrial respiration was subtracted to obtain basal, coupled, uncoupled (proton leak) and maximal mitochondrial respiration.

ATP levels were quantified by the ATP Determination Kit (Invitrogen). Cells were washed 48 hours post-transfection and preincubated for 1 hour in KRB (129 mM NaCl, 4.8 mM KCl, 1 mM CaCl_2_, 5 mM NaHCO_3_, 1.2 mM KH_2_PO4, 1.2 mM MgSO_4_, 10 mM HEPES, 0.1% BSA, pH 7.4) containing low glucose (2 mM). Subsequently, cells were incubated for 1 hour under either basal (2 mM glucose) or stimulatory (25 mM glucose) conditions. To inhibit mitochondrial pyruvate uptake, cells were treated with 2 µM UK-5099, a specific inhibitor of the mitochondrial pyruvate carrier. To inhibit the entry of fatty acids bound to albumin present in the assay buffer into mitochondria, 100 µM etomoxir was used. Glucose metabolism was blocked by the addition of 2-deoxyglucose. All compounds were purchased from Sigma–Aldrich. Finally, culture supernatant was quickly discarded and cells were lysed in Passive lysis buffer (Promega). Intracellular ATP concentrations were quantified in cell lysates using a bioluminescence-based ATP Determination Kit (Invitrogen), following the manufacturer’s protocol. ATP levels were normalized to protein content determined by Pierce™ BCA Protein Assay Kits (ThermoFisher Scientific). Luminescence and absorbance were determined by SynergyMx Plate microplate reader (Biotek).

Reactive oxygen species were measured using a non-lytic assay of ROS-Glo™ H2O2 Assay kit (Promega).

#### Cell viability

Inflammation was induced with IL1β (10 ng/ml) and IFNγ (100 ng/ml) for 24 hours. Cell viability was measured using the AlamarBlue™ HS reagent (Invitrogen).

#### Transmission electron microscopy

MIN6 cells 48h post-transfection were mildly trypsinized with 0.05% Trypsin-EDTA for 2 min at room temperature (RT) and subsequently detached by tapping the plate in order to minimize changes in cell morphology. Cell suspension was centrifuged at 1000 rpm/5 min/RT and the cell pellet was fixed for 2 hours in 2% glutaraldehyde in cacodylate buffer and post-fixed in 1% osmium tetroxide (OsO₄) in the same buffer. After washing, samples were contrasted with 1% aqueous uranyl acetate. Dehydration was carried out through a graded ethanol series followed by propylene oxide, after which samples were infiltrated with a 1:1 mixture of propylene oxide and Durcupan resin. Polymerization was performed in pure Durcupan at 60 °C for 72 hours. Ultrathin sections (60 nm) were prepared using a PowerTome MT-XL ultramicrotome (RMC/Sorvall, USA), contrasted with lead citrate and imaged using a JEM 1200 transmission electron microscope (JEOL, Japan) operated at 80 kV. Images were acquired with a CCD camera (Dual Vision 300W, Gatan, USA).

#### Determination of intracellular calcium

Cells were loaded with 1 μM Fluo-3-AM probe (Sigma-Aldrich) for 1 hour at 37 °C, 5 % CO_2_ in serum-free low glucose medium in the presence/absence of EDTA. After washing with PBS, fluorescence was measured at λex 485 nm and λem 520 nm.

#### Protein extraction and western blot

Cells were lysed in RIPA buffer (50 mM Tris-HCl pH 7.4, 150 mM NaCl, 2 mM EDTA, 1.0% Triton X-100, 0.5% sodium deoxycholate) supplemented with protease and phosphatase inhibitors (Roche). Lysates were probed with primary antibodies listed in Supplementary Table 2. Signals of IRDye-labeled secondary antibodies were detected by Odyssey Infrared Imaging System (LiCor). Densitometric analysis of immunoreactive bands was conducted in ImageJ (version 1.53e, NIH).

#### Statistical analysis

Sample sizes were guided by previous studies using comparable experimental designs; individual datapoints are shown in figures and numbers are listed in figure legends. All animals were analysed, and blinding was not applied. Cell culture studies used 2-3 technical replicates for mRNA/protein and 5-6 for respiration/lipolysis, and were reproduced 2-4 times. Data are presented as mean ± SEM. Statistical significance was determined using GraphPad Prism 8 by unpaired two-tailed Student’s t-test to compare two groups; ordinary two-way ANOVA with Sidak’s or Tukeýs multiple comparison test to analyse the effect of two factors; or two-way ANOVA with repeated measurements with Sidak post-hoc test to analyse parameters measured over time. The use of each test is indicated in the figure legends. Statistical differences are indicated as * p < 0.05, ** p < 0.01 and *** p < 0.001.

## Results

### GPR180 dysfunction attenuates glucose-stimulated insulin secretion

Global deletion of GPR180 receptor resulted in impaired glucose tolerance in chow-fed mice (Fig. 1A), accompanied by elevated random fed glycaemia (Fig. 1B) but significantly reduced plasma insulin level (Fig. 1C). Insulin tolerance remained unchanged (Fig. 1D), indicating that the phenotype arises from altered β-cell function rather than impaired peripheral insulin sensitivity.

**Figure 1.**
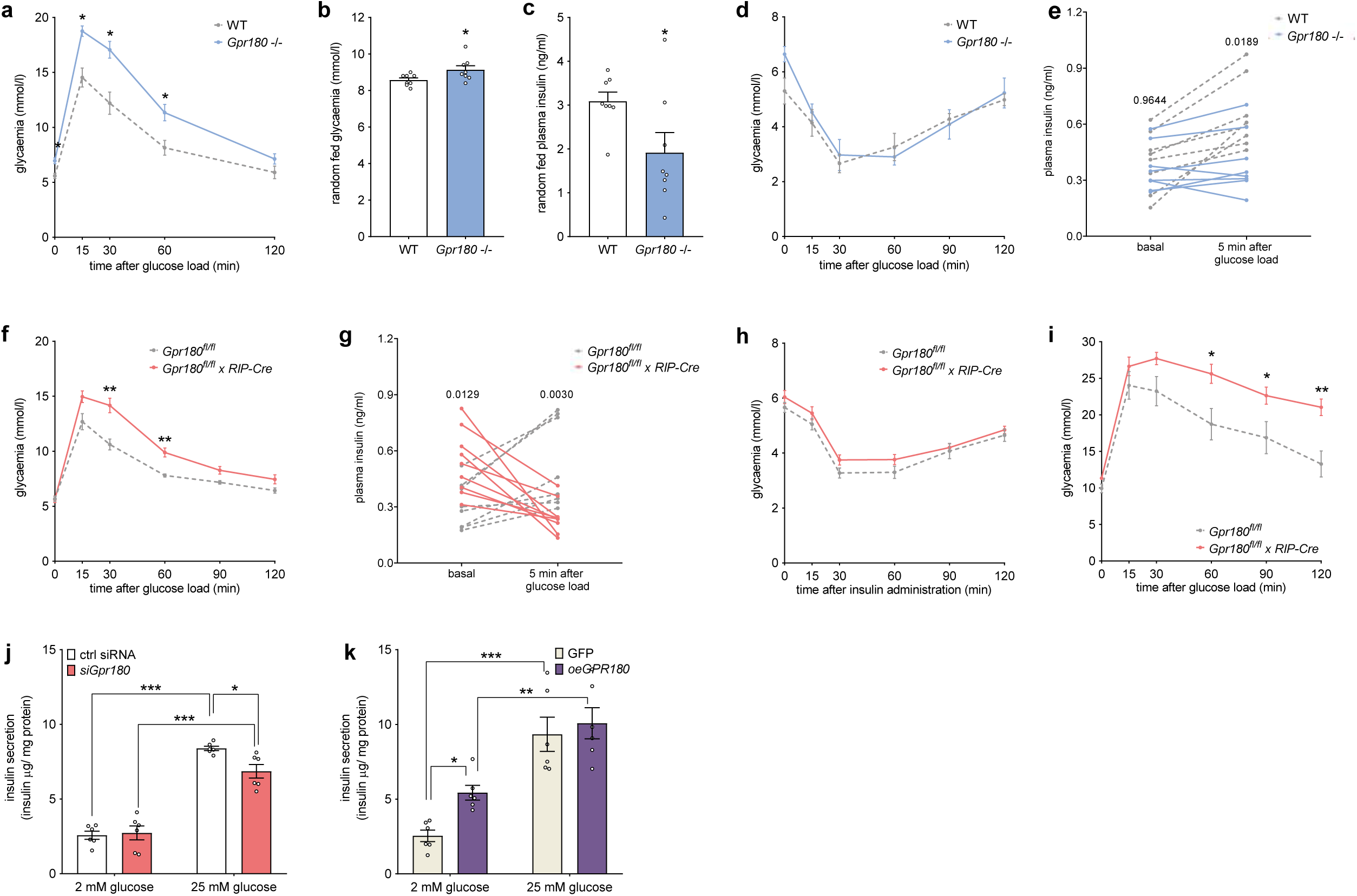
GPR180 deficiency impairs glucose-stimulated insulin secretion in a β- cell-autonomous manner. (A) Glucose tolerance test (GTT) (n = 5 for WT, n = 7 for *Gpr180* global knockout), (B) random fed blood glucose levels, (C) random fed plasma insulin concentrations, (D) insulin tolerance test (ITT) (n = 5 for WT, n = 7 for *Gpr180* global knockout) and (E) first-phase insulin secretion in response to glucose administration in chow-fed global *Gpr180* knockout and wild-type mice. (F) GTT (n = 7 for *Gpr180^fl/fl^*, n = 9 for *Gpr180^fl/fl^ x RIP-Cre*), (G) first-phase insulin secretion in response to glucose administration, (H) ITT (n = 8 for both genotypes) in chow-fed β-cell-specific *Gpr180* knockout mice and Cre-negative controls. (I) GTT in high-fat diet (HFD)-fed β-cell-specific *Gpr180* knockout mice and Cre-negative controls (n = 8 for both genotypes). Glucose-stimulated insulin secretion in MIN6 cells following (J) *Gpr180* silencing and (K) *GPR180* overexpression. Data are presented as mean ± SEM. Statistical significance was determined using an unpaired two-tailed t-test (B, C), ordinary two-way ANOVA with subsequent Tukey’s multiple comparison test (E, G, J, K), or two-way ANOVA with repeated measures followed by Sidak’s post hoc test when appropriate (A, D, F, H, I). * p < 0.05; ** p < 0.01; *** p < 0.001. Gpr180 G-protein coupled receptor 180; RIP rat insulin promoter; WT wild-type.

To address this, we measured first-phase insulin secretion in response to glucose administration. Global *Gpr180* knockout mice showed a blunted insulin response (Fig. 1E and supplementary Fig. 1A), revealing a defect in GSIS.

To determine whether this effect is β-cell-intrinsic, we generated a β-cell-specific *Gpr180* knockout model (b*Gpr180*-KO) (Supplementary Fig. 1B). These mice phenocopied the global knockout, displaying impaired glucose tolerance (Fig. 1F) and defective first-phase insulin secretion (Fig. 1G and Supplementary Fig. 1C) without differences in body weight (Supplementary Fig. 1D) or insulin tolerance (Fig. 1H). When subjected to a high-fat diet, b*Gpr180*-KO mice showed exacerbated glucose intolerance and worsened diabetic phenotype (Fig. 1I), again independent of body weight changes (Supplementary Fig. 1E).

To support these findings, MIN6 β-cells were used for *in vitro* gain- and loss-of-function studies as a controlled model to investigate the cellular basis of GPR180 function. *Gpr180* silencing (Supplementary Fig. 1F) significantly reduced GSIS (Fig. 1J, Supplementary Fig. 1G), whereas overexpression (Supplementary Fig. 1H) enhanced basal insulin secretion (Fig. 1K, Supplementary Fig. 1I).

Together, these findings demonstrate that GPR180 is required for efficient GSIS in a β-cell-autonomous manner.

### GPR180 Controls β-Cell Fuel Processing and Metabolic Flux

To investigate the molecular basis of impaired insulin secretion following loss of GPR180, we first performed bulk RNA-seq on MIN6 cells with silenced *Gpr180*. Transcriptomic alterations were modest, with only 62 DEGs (Supplementary data 1). Gene ontology (GO) analysis revealed enrichment in pathways related to vesicular transport, insulin secretion, and cellular responses to TGFβ (Fig. 2A), while expression of insulin genes and key transcription factors regulating insulin synthesis remained unchanged (Fig. 2B). Among the downregulated genes, we noted reduced expression of *Serp1* and *Syt11*, involved in ER function and vesicle dynamics, respectively, whereas Slc16a1 was not considered further due to its known aberrant expression in MIN6 cells and its minimal expression in mature β-cells under physiological conditions [17] (Fig. 2C). These data suggest that impaired insulin secretion is unlikely to result from altered hormone production or transcriptional reprogramming.

**Figure 2.**
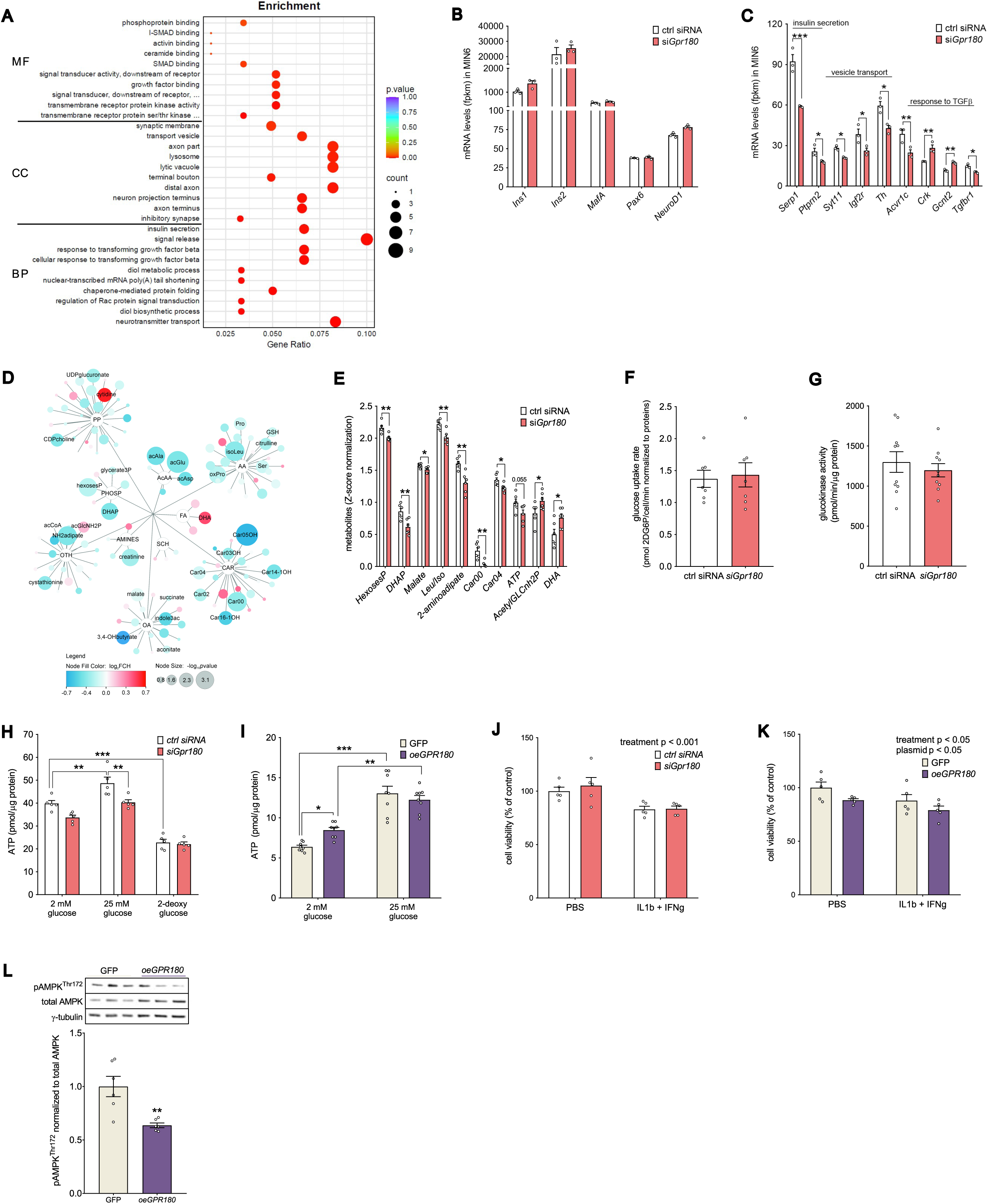
GPR180 affects insulin secretion by modulating mitochondrial glucose metabolism. (A) Gene ontology enrichment analysis of differentially expressed genes from RNA-sequenced MIN6 cells following *Gpr180* silencing, showing the top 10 molecular functions, cellular components, and biological processes. Expression of (B) insulin genes and key β-cell transcription factors and (C) genes implicated in insulin secretion, vesicle transport, and TGFβ response in MIN6 cells with attenuated *Gpr180* expression. (D) Cytoscape network visualization of altered metabolites. Node size reflects the −log(p-value) and node color indicates the fold change (red = increased, blue = decreased in *Gpr180*-deficient cells vs. controls). Metabolites are clustered by biochemical class, and only those that are statistically significant and/or discussed in the text are shown. (E) Relative levels of selected metabolites related to glycolysis (hexose phosphates, DHAP), TCA cycle (malate), amino acids (Iso/Leu, 2-aminoadipate), acylcarnitines (Car00, Car04), and Hexosamine/lipid-related metabolites (AcetylGLCnh2P, DHA). (F) Glucose uptake and (G) glucokinase activity in MIN6 following *Gpr180* knockdown. Cellular ATP levels in response to glucose stimulation of MIN6 following (H) *Gpr180* silencing and (I) GPR180 overexpression. Cell viability under basal or cytokine-induced stress in MIN6 with (J) *Gpr180* silencing and (K) *GPR180* overexpression. (L) AMPK phosphorylation in MIN6 overexpressing GPR180 was determined by Western blot. Data are presented as mean ± SEM. Statistical analysis was performed by unpaired t-test (B, C, E – G, L) or ordinary two-way ANOVA with Sidak’s post-hoc analysis (H - K). * p < 0.05; ** p < 0.01; *** p < 0.001. AA amino acids; AcAA acetylated amino acids; acAla acetylalanine; acAsp acetylaspartate; acCoA acetyl-CoA; acGlu acetylglutamate; AcetylGLCnh2P N-acetylglucosamine-6-phosphate; *Acvr1c* activin A receptor, type IC; AMINES metabolites containing an amino group; ATP adenosine triphosphate; BP biological process; Car acylcarnitines; Car00 free carnitine; Car02 acetylcarnitine; Car03OH 3-hydroxypropionylcarnitine; Car04 butyrylcarnitine; Car05OH 3-hydroxyvalerylcarnitine; Car14-1OH 3-hydroxytetradecenoylcarnitine; Car16-1OH 3-hydroxyhexadecenoylcarnitine; CC cellular component; *Crk* v-crk avian sarcoma virus CT10 oncogene homolog; DHA docosahexaenoic acid; DHAP dihydroxyacetone phosphate; FA fatty acids (non-esterified); *Gcnt2* glucosaminyl (N-acetyl) transferase 2; *Gpr180* G-protein coupled receptor 180; GSH glutathione; HesosesP hexose phosphates; IFNγ interferon gamma; *Igf2r* insulin-like growth factor 2 receptor; IL1β interleukin 1 beta; indole3ac indole-3-acetate; *Ins1* Insulin 1; *Ins2* Insulin 2; Leu/Iso Leucine/Isoleucine; *MafA* MAF bZIP transcription factor A; MF molecular function; *NeuroD1* neurogenic differentiation 1; OA organic acids; OTH others; *Pax6* paired box 6; PBS phosphate buffered saline; PHOSP metabolites containing a phosphate group; PP purines/pyrimidines; *Ptprn2* protein tyrosine phosphatase receptor type N polypeptide 2; RFU relative fluorescence units; SCH saccharides; *Serp1* stress-associated endoplasmic reticulum protein 1; *Slc16a1* solute carrier family 16 (monocarboxylic acid transporters) member 1; *Syt11* synaptotagmin XI; *Tgfbr1* transforming growth factor, β receptor I; *Th* tyrosine hydroxylase.

To address this, we performed metabolomic profiling of MIN6 cells after *Gpr180* silencing. Targeted analysis revealed reductions in multiple metabolite classes (Fig. 2D), including amino acids (e.g., serine, proline, leucine/isoleucine), acetylated amino acids (acetylglutamate, acetylalanine, acetylaspartate), TCA-cycle–linked intermediates (malate), glycolytic and glycerolipid pathway components (hexose phosphates, dihydroxyacetone phosphate, CDP-choline), and short acylcarnitines (Fig. 2E). ATP content was lower in *Gpr180*-deficient cells, although this difference did not reach statistical significance (p = 0.055), suggesting reduced cellular energy production. Conversely, levels of docosahexaenoic acid and the hexosamine-pathway metabolite N-acetylglucosamine-phosphate were increased, indicating altered lipid and amino sugar metabolism. These changes are consistent with alterations in central carbon metabolism and may reflect a shift toward lipid remodeling and the hexosamine biosynthetic pathway under reduced mitochondrial oxidation. Although several intermediates (3-phosphoglycerate, acetyl-CoA, aconitate and succinate) did not reach statistical significance individually, their coordinated downward trends in *Gpr180*-deficient cells, together with significant changes in related metabolites, suggest a pathway-level alteration in metabolic flux that may contribute to impaired fuel processing in β-cells. In line with this, metabolic pathway analysis using MetaboAnalyst identified enrichment of pathways primarily related to amino acid, glycerophospholipid, inositol phosphate and glutathione metabolism (Supplementary Fig. 2A, B).

Since early steps in glucose handling could contribute to these alterations, we assessed glucose uptake and glucokinase activity. Both remained unchanged after *Gpr180* silencing (Fig. 2F, G), indicating that glucose sensing upstream of glycolysis is intact. Instead, *Gpr180*-deficient cells exhibited a clear defect in glucose-stimulated ATP production (Fig. 2H). No difference was observed when cells were exposed to 2-deoxyglucose, confirming that the impairment most likely arises from metabolic processing rather than glucose entry or phosphorylation. To exclude an increased reliance on fatty acid oxidation as a confounding factor, ATP measurements were repeated in the presence of the CPT1a inhibitor etomoxir. The reduction in glucose-stimulated ATP production persisted (Supplementary Fig. 2C), demonstrating that the effect is independent of shifts in lipid oxidation. Conversely, *GPR180* overexpression increased basal ATP levels (Fig. 2I), reflecting the elevated basal insulin secretion observed earlier. Cell viability assessed by resazurin reduction was unchanged following *Gpr180* knockdown (Fig. 2J), whereas a modest decrease in viability with *GPR180* overexpression (Fig. 2K) likely reflects plasmid-associated stress rather than true metabolic impairment. In line with increased ATP availability, AMPK phosphorylation was reduced upon *GPR180* overexpression (Fig. 2L), suggesting altered cellular energy status.

These findings position GPR180 as a determinant of β-cell metabolic function associated with insulin secretion, acting downstream of glucose uptake and phosphorylation to facilitate efficient ATP generation, which is essential for triggering insulin granule exocytosis during GSIS.

### GPR180 Regulates Mitochondrial Function and Microenvironment

Our metabolomic analyses revealed coordinated reductions in glycolytic and TCA cycle–related intermediates, as well as diminished levels of acylcarnitines and amino acids, suggesting compromised mitochondrial substrate entry and diminished anaplerotic support of the TCA cycle. Together with lower cellular ATP levels, this metabolic profile indicates impaired mitochondrial substrate utilization and limited metabolic flexibility. This prompted us to investigate mitochondria as a potential basis for the impaired energetic and secretory phenotype in *Gpr180*-deficient β-cells.

To determine whether altered pyruvate-derived anaplerosis contributes to reduced ATP generation, we pharmacologically inhibited the mitochondrial pyruvate carrier (MPC). This inhibition of pyruvate entry to mitochondria abolished the difference in glucose-stimulated ATP production between control and *Gpr180*-silenced cells (Fig. 3A), indicating that pyruvate-driven anaplerotic flux is already functionally constrained in the absence of GPR180. We next asked whether impaired mitochondrial metabolism reflected changes in mitochondrial abundance. Neither mitochondrial DNA copy number, OXPHOS complex abundance, nor mitochondrial mass assessed by MitoTracker Green staining was altered by *Gpr180* knockdown or overexpression (Fig. 3C–F), demonstrating that GPR180 does not regulate mitochondrial quantity. In contrast, mitochondrial quality and function were markedly impaired. *Gpr180*-deficient MIN6 cells exhibited a reduced JC-1 red/green fluorescence ratio, demonstrating diminished mitochondrial membrane polarization (Fig. 3G). Seahorse respirometry revealed overall suppression of mitochondrial respiration, including lower basal, ATP-linked, and maximal oxygen consumption (Fig. 3H). The decline in basal and coupled respiration aligns with mitochondrial depolarization and decreased ATP levels, indicating a functional impairment of mitochondrial activity. However, *GPR180* overexpression did not significantly alter the mitochondrial oxygen consumption rate (Fig. 3I), suggesting that endogenous GPR180 levels are required to maintain mitochondrial function, whereas supra-physiological increases do not further enhance respiration. Supporting impaired mitochondrial and metabolic function, *Gpr180*-deficient cells generated significantly higher levels of reactive oxygen species (Fig. 3J). Overexpression of *GPR180* did not significantly affect ROS levels (Fig. 3K), again pointing to a requirement for physiological, but not excessive, GPR180 activity to preserve mitochondrial homeostasis.

**Figure 3.**
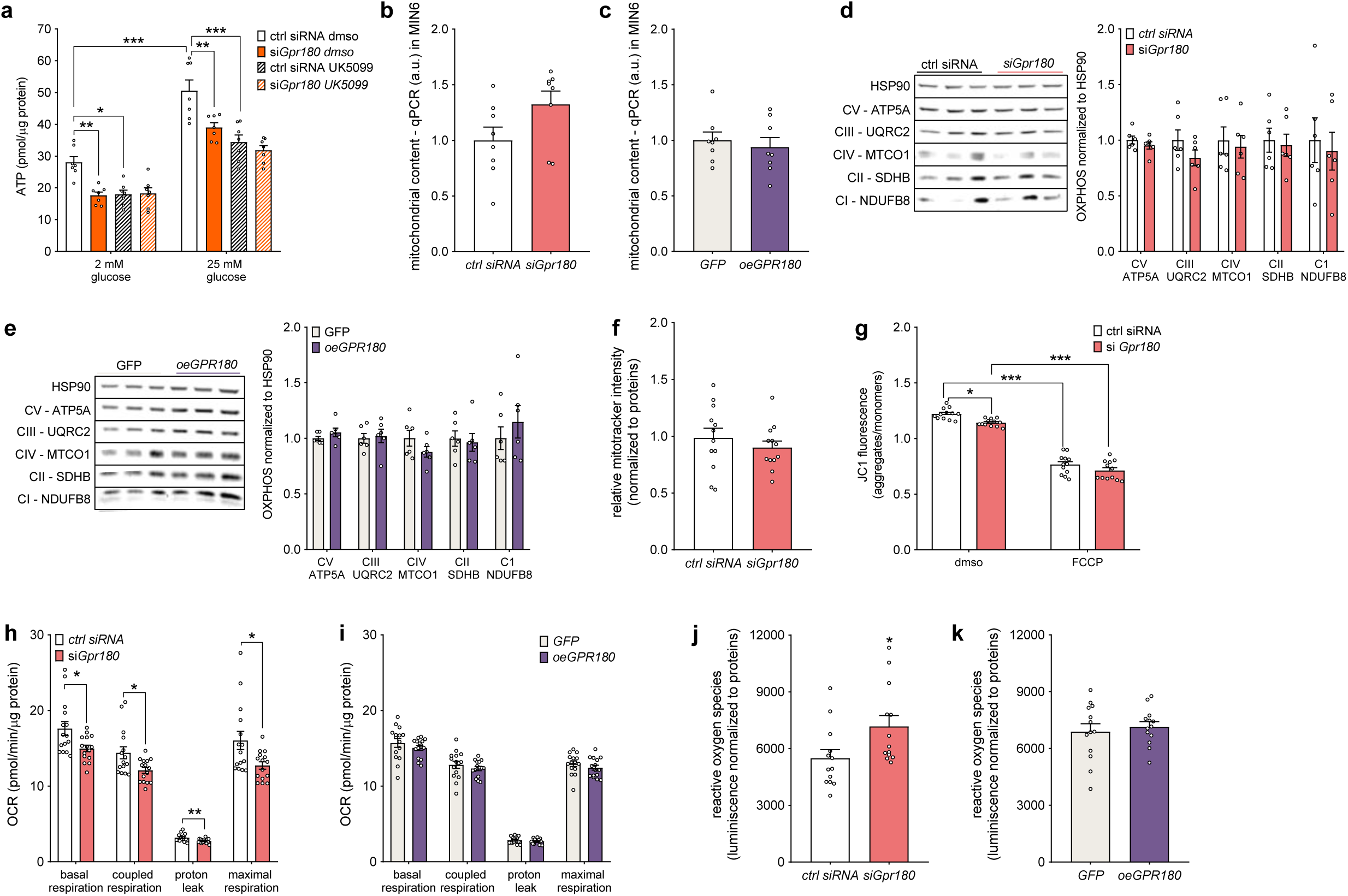
GPR180 alters mitochondrial function. (A) ATP levels in control MIN6 cells and following *Gpr180* silencing in response to inhibition of mitochondrial pyruvate carrier with UK-5099. Mitochondrial DNA copy number in MIN6 following (B) *Gpr180* knockdown and (C) overexpression. Western blot analysis of OXPHOS complex protein levels following (D) *Gpr180* silencing and (E) overexpression. (F) MitoTracker Green staining (mitochondrial mass) and (G) JC-1 staining (mitochondrial membrane potential) in *Gpr180*-silenced MIN6 cells. Cellular respiration determined by Seahorse extracellular flux analysis (oxygen consumption rate, OCR) in MIN6 cells following (H) *Gpr180* knockdown and (I) overexpression. Reactive oxygen species in MIN6 following (J) *Gpr180* knockdown and (K) overexpression. Data are shown as mean ± SEM. Statistical analysis was performed by an unpaired two-tailed t-test (B – F, H - K) or ordinary two-way ANOVA with subsequent Sidak’s post-hoc test (A, G). * p < 0.05; ** p < 0.01; *** p < 0.001. ATP adenosine triphosphate, ATP5A ATP synthase F1 subunit α, DMSO Dimethyl Sulfoxide, *Gpr180* G-protein coupled receptor 180, HSP90 heat shock protein 90, MTCO1 mitochondrially encoded cytochrome c oxidase I, NDUFB8 NADH dehydrogenase [ubiquinone] 1 β subcomplex subunit 8, OCR oxygen consumption rate, SDHB Succinate Dehydrogenase Complex Iron Sulfur Subunit B, UQRC2 Ubiquinol-Cytochrome C Reductase Core Protein 2.

Given the observed mitochondrial functional defects, we next asked whether these are accompanied by changes in subcellular organization that could influence mitochondrial performance. To address this, we performed ultrastructural analysis of MIN6 cells by transmission electron microscopy to examine structural basis for impaired mitochondrial function. Loss of *Gpr180* was associated with altered organization of the endoplasmic reticulum (ER) network (Fig. 4A, B), and a reduced presence of ER reticular structures in proximity to mitochondria, such that mitochondria appeared spatially decoupled from the ER (Fig. 4C, D). Consistent with ER alterations, *Gpr180*-deficient β-cells exhibited lower basal cytosolic Ca²⁺ levels under low-glucose conditions, an effect abolished by extracellular Ca²⁺ chelation, indicating altered intracellular Ca²⁺ handling (Fig. 4E). In contrast, cells overexpressing *GPR180* exhibited a more extensive and organized ER network, enlarged Golgi structures, and mitochondria closely juxtaposed to ER membranes (Supplementary Fig. 3A–D). These alterations in subcellular architecture indicate that GPR180 is involved in maintaining the mitochondrial microenvironment required for metabolic coupling, as ER–mitochondria proximity has been implicated in the regulation of Ca²⁺ exchange and insulin secretion in β-cells [18]. Our findings indicate that GPR180 contributes to maintaining mitochondrial integrity and function, including membrane potential and respiration, as well as appropriate inter-organelle organization that supports β-cell energy metabolism and insulin release, thereby providing mechanistic insight into how loss of GPR180 impairs β-cell function. These effects occur without changes in mitochondrial mass, indicating that GPR180 primarily affects mitochondrial quality and microenvironment rather than abundance.

**Figure 4.**
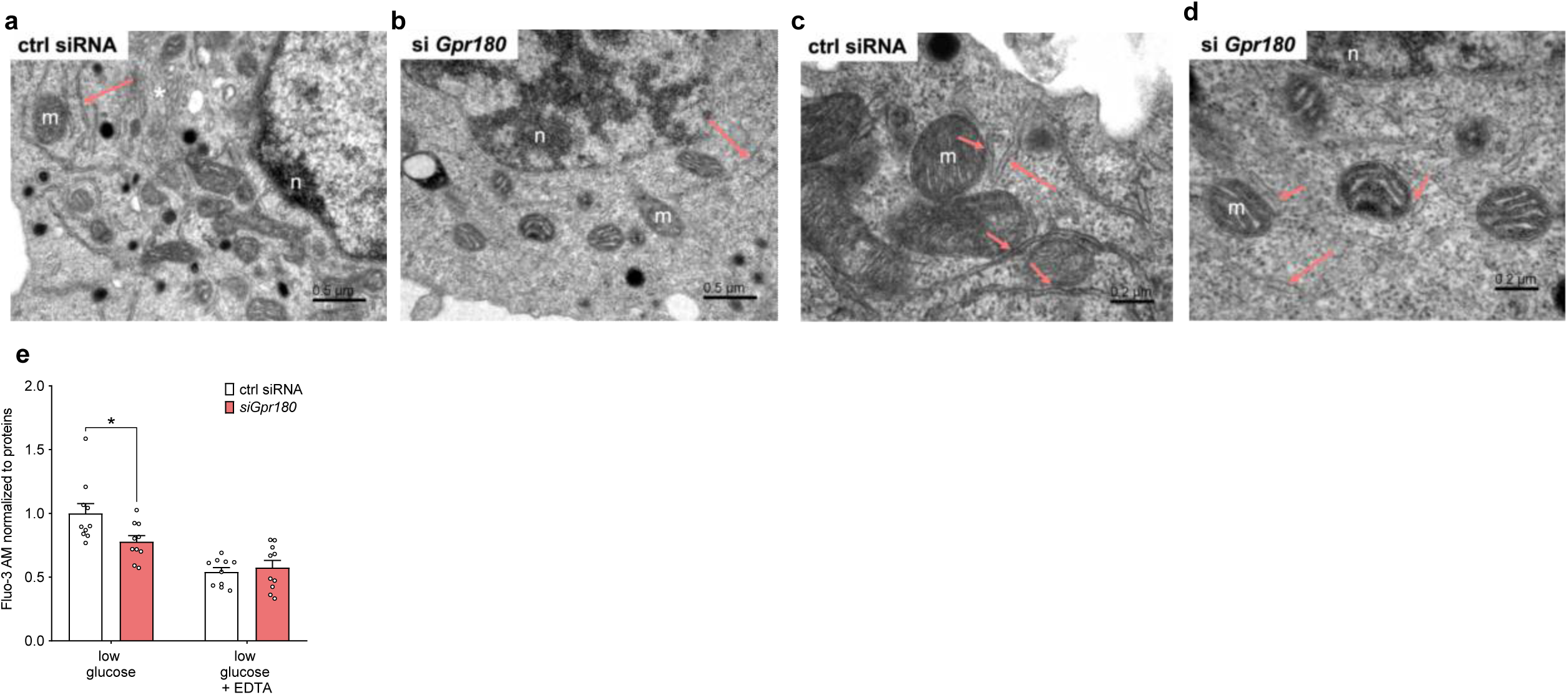
GPR180 alters cellular morphology and mitochondrial microenvironment. View of the area of the cell with transfected (A) control siRNA and (B) siRNA targeting *Gpr180*. Higher-magnification views of the mitochondrial environment in (C) control cells and (D) cells following *Gpr180* silencing. (A, B) scale bar 500 nm and (C, D) scale bar 200 nm; m - mitochondria, n - nucleus, asterisk - Golgi apparatus, long arrows - endoplasmic reticulum, and short arrows - endoplasmic reticulum in proximity to the outer mitochondrial membrane. (E) Intracellular calcium levels in control MIN6 cells and following *Gpr180* silencing under low-glucose conditions in the presence or absence of EDTA chelator. *Gpr180* G-protein coupled receptor 180. Data are shown as mean ± SEM. Statistical analysis was performed by ordinary two-way ANOVA with subsequent Sidak’s post-hoc test (E). * p < 0.05.

### GPR180 Is Required for Metabolic Fitness and Endocrine Identity in Pancreatic Islets

To assess the physiological relevance of our *in vitro* findings, we performed a comprehensive analysis of pancreatic architecture in b*Gpr180*-KO mice. Given the observed effects of GPR180 on β-cell function and mitochondrial activity, we asked whether these alterations are evident in the endocrine pancreas *in vivo*.

Histomorphometric analysis revealed no significant differences in pancreatic mass (Fig. 5A) or overall pancreatic architecture (Fig. 5B). Immunohistochemical staining for insulin further demonstrated that the number and size of Langerhans islets, as well as total β-cell mass, were preserved in b*Gpr180*-KO mice (Fig. 5C-F). Despite the intact islet structure, total pancreatic insulin content was significantly lower in bGpr180-KO (Fig. 5G), suggesting a functional rather than structural deficit.

**Figure 5.**
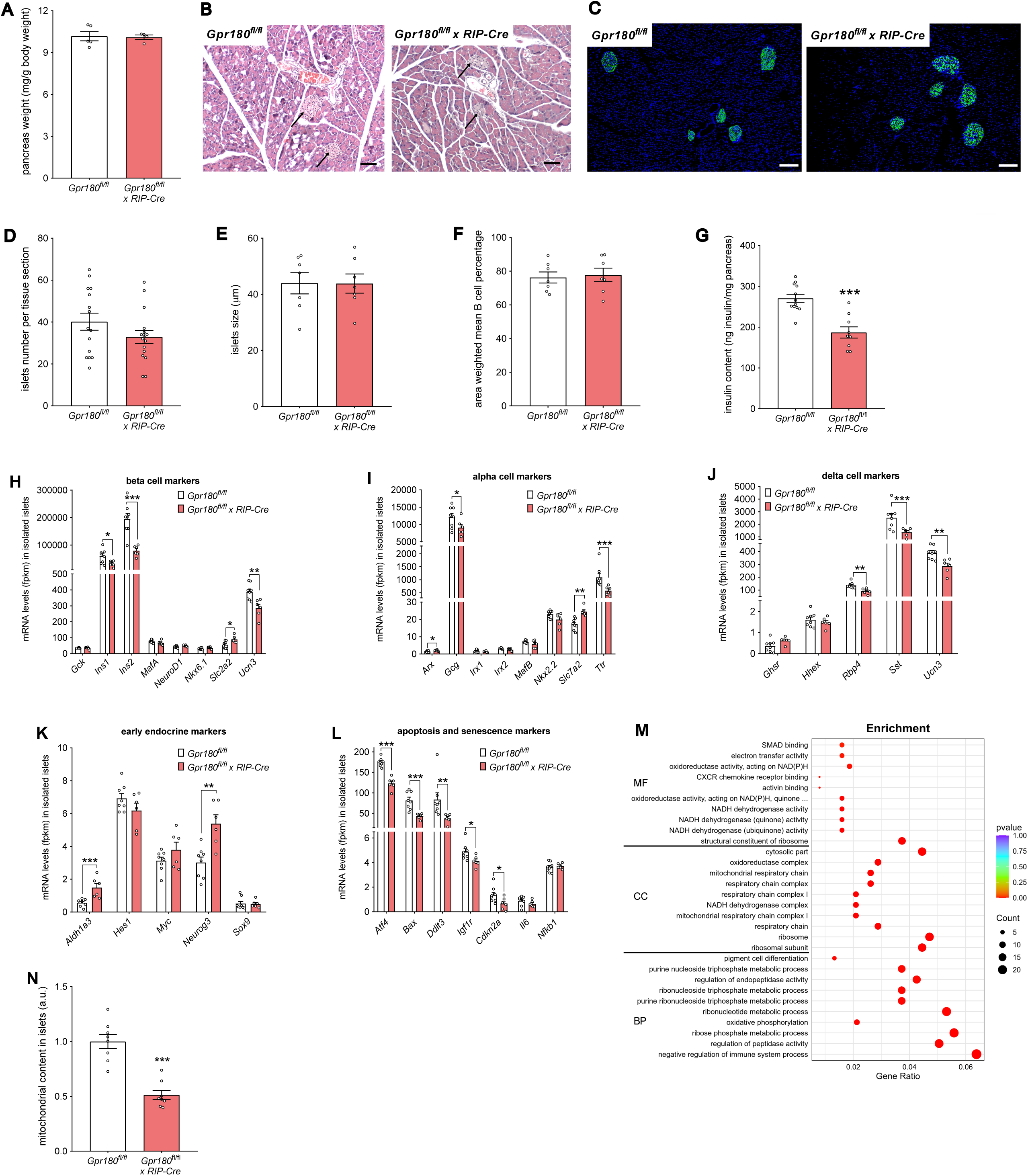
Loss of GPR180 alters mitochondrial abundance and endocrine cell identity in pancreatic islets. (A) Pancreatic mass in β-cell-specific *Gpr180* knockout mice and controls. (B) Hematoxylin and eosin staining of pancreatic sections(magnification 10x, scale bar 50 µm, arrows indicating islets of Langerhans) (C) Representative images of pancreas sections of control and β-cell-specific *Gpr180* knockout mice stained with anti-insulin antibody (green) and DAPI (blue) (magnification 5x, scale bar 50 µm). Quantification of (D) islet number per tissue section, (E) islet size, and (F) β-cell mass in controls and mice lacking *Gpr180* specifically in β-cells. (G) Total pancreatic insulin content in β-cell-specific *Gpr180*-deficient mice compared to controls. Expression level of (H) β-cell-, (I) α-cell-, (J) δ-cell- specific markers based on RNA-seq of isolated pancreatic islets from control and β-cell-specific *Gpr180* knockout mice. Expression of (K) endocrine progenitor markers, (L) markers of cellular stress, apoptosis, and senescence in *Gpr180*-deficient and control islets. (M) Gene ontology enrichment analysis of differentially expressed genes from RNA-sequenced isolated Langerhans islets from the pancreas of control and β-cell-specific Gpr180 knockout mice, showing the top 10 molecular functions, cellular components, and biological processes. (N) Quantification of mitochondrial content in isolated pancreatic islets from β-cell-specific *Gpr180-ablated* mice and control mice. Data are presented as mean ± SEM. Statistical analysis was performed using an unpaired two-tailed t-test. * p < 0.05; ** p < 0.01; *** p < 0.001. *Aldh1a3* aldehyde dehydrogenase family 1 subfamily A3, *Arx* aristaless related homeobox, *Atf4* activating transcription factor 4, *Bax* BCL2-associated X protein, *Cdkn2a* cyclin dependent kinase inhibitor 2A, *Ddit3* DNA-damage inducible transcript 3, *Gcg* glucagon, *Gck* glucokinase, *Ghsr* growth hormone secretagogue receptor, *Gpr180* G-protein coupled receptor 180, *Hes1* hes family bHLH transcription factor 1, *Hhex* hematopoietically expressed homeobox, *Igf1r* insulin-like growth factor I receptor, *Il6* interleukin 6, *Ins1* Insulin 1, *Ins2* Insulin 2, *Irx1* Iroquois homeobox 1, *Irx2* Iroquois homeobox 2, *MafA* MAF bZIP transcription factor A, *MafB* MAF bZIP transcription factor B, *Myc* myelocytomatosis oncogene, *NeuroD1* neurogenic differentiation 1, *Neurog3* neurogenin 3, *Nfkb1* nuclear factor of kappa light polypeptide gene enhancer in B cells 1, *Nkx2.2* NK2 homeobox 2, *Nkx6.1* NK6 homeobox 1, *Rbp4* retinol binding protein 4, RIP rat insulin promoter, *Slc2a2* solute carrier family 2 (facilitated glucose transporter), member 2, *Slc7a2* solute carrier family 7 (cationic amino acid transporter, y+ system), member 2, *Sox9* SRY (sex determining region Y)-box 9, *Sst* somatostatin, *Ttr* transthyretin, *Ucn3* urocortin 3.

To explore the molecular basis of this phenotype, we performed bulk RNA-seq on isolated islets from control and b*Gpr180*-KO mice. Transcriptomic profiling revealed downregulation of β-cell–enriched gene signatures (Fig. 5H), accompanied by reduced expression of α-cell (Fig. 5I) and δ-cell–specific markers (Fig. 5J) [19]. Conversely, several markers associated with endocrine progenitor and less differentiated states were upregulated (Fig. 5K), indicating a shift away from mature endocrine cell identity. Notably, this transcriptional reprogramming occurred in the absence of an overt stress response, as genes associated with apoptosis, cellular senescence, and inflammation were not induced and, in some cases, were decreased (Fig. 5L). GO analysis of DEGs identified enrichment of mitochondrial respiratory chain components and oxidative phosphorylation as one of the key cellular components and biological functions dysregulated in b*Gpr180*-KO islets (Fig. 5M, Supplementary data 2), indicating diminished mitochondrial metabolic capacity under GPR180 deficiency *in vivo*. In line with this, we detected a substantial reduction in mitochondrial content in the islets of b*Gpr180*-KO mice (Fig. 5N), suggesting long-term mitochondrial remodeling likely constraining ATP-generating capacity, reflecting the compromised mitochondrial integrity observed in our *in vitro* model. These *in vivo* data support a role for GPR180 in maintaining mitochondrial metabolism and β-cell identity in pancreatic islets.

## Discussion

Diabetes is characterized by progressive dysregulation of glucose homeostasis, largely driven by impaired β-cell function and loss of insulin secretory capacity. Despite numerous therapeutic options, optimizing diabetes management remains a major challenge. In this study, we identify GPR180 as a previously unrecognized regulator of pancreatic β-cell function that is linked to mitochondrial metabolism and insulin secretion. Using complementary *in vivo* and *in vitro* models, we show that loss of GPR180 impairs GSIS and glucose tolerance through β-cell-autonomous mechanisms. This phenotype is characterized by reduced mitochondrial respiration and ATP production, indicating compromised bioenergetic capacity. Metabolomic profiling revealed decreased levels of glycolytic and TCA cycle intermediates, along with lower acylcarnitines and amino acids, indicating reduced metabolic flux and impaired mitochondrial substrate utilization *in vitro* following *Gpr180* silencing. These changes indicate an altered allocation of glucose- and lipid-derived substrates away from mitochondrial oxidation, which may contribute to diminished anaplerotic support of the TCA cycle, the processes previously linked to defective insulin secretion and β-cell failure in diabetes [2, 6, 20]. In parallel, ultrastructural ER remodeling and altered intracellular Ca²⁺ handling suggest disruption of the mitochondrial microenvironment, potentially weakening the metabolic coupling required for efficient GSIS, as reported previously [18, 21]. Together, these findings support a model in which early metabolic alterations precede transcriptional remodelling, with reduced expression of oxidative phosphorylation genes and mitochondrial content in b*Gpr180*-KO islets likely reflecting long-term adaptation to sustained metabolic stress. Notably, these changes were accompanied by alterations in endocrine cell identity. This is consistent with emerging evidence that mitochondrial dysfunction is closely linked to loss of β-cell maturity and dedifferentiation, as highlighted in recent studies [22]. In this context, mitochondrial integrity appears to be important for maintaining β-cell identity, and its disruption may contribute to functional decline under metabolic stress. Collectively, these findings position GPR180 as a modulator of β-cell metabolic competence, linking organelle function, substrate utilization, and secretory capacity. A major strength of this study is the integration of multiple experimental systems and methodologies. The use of both whole-body and β-cell-specific knockout [23] models together with gain- and loss-of-function experiments in MIN6 cells [24, 25], alongside detailed analyses of transcriptome, metabolome, mitochondrial bioenergetics, and ultrastructure, allowed us to investigate the cellular basis of GPR180 function in β-cells across complementary systems.

Previous studies have linked GPR180 to adipose and hepatic metabolism [7–9, 26]. Here, we extend these observations by demonstrating a role of GPR180 in the endocrine pancreas. Importantly, the mitochondrial dysfunction and altered endocrine cell identity observed in *Gpr180*-deficient islets align with previous studies linking impaired mitochondrial metabolism to β-cell dedifferentiation and diabetes progression [22, 27–29], reinforcing the physiological relevance of our findings. We previously identified GPR180 as a component of the canonical TGFβ signalling machinery in various human cell lines [7]. However, we did not observe clear evidence for involvement of the canonical TGFβ pathway in murine β-cells. This discrepancy may reflect cell–type–specific or species-dependent differences, or limitations of the MIN6 model. Instead, transcriptomic and GO analyses of both MIN6 cells and isolated islets with ablated GPR180 suggest engagement of alternative branches of the non-canonical TGFβ superfamily, such as activin receptor–associated pathways. Together with recent studies indicating the pleiotropic nature of GPR180 [9, 26, 30], these findings support the notion that its downstream signalling is context-dependent.

Our findings have clinically relevant implications. Preservation of β-cell function remains a therapeutic goal in both type 1 and type 2 diabetes, and mitochondrial dysfunction is recognized as a contributor to β-cell failure [28, 31]. The plasma membrane localization of GPR180 makes it an attractive, potentially druggable target. Modulation of GPR180 activity in β-cells may help preserve mitochondrial function and insulin secretion under metabolic stress, although this requires further validation. Furthermore, genome-wide association data from the Diabetes Knowledge Portal [32] have identified an association of a single nucleotide polymorphism (SNP) in the *GPR180* gene locus (rs6492721 T>C) with variation in HbA_1c_ levels (p=1.183e-14). This provides independent support for a functional role of GPR180 in regulating glucose homeostasis, consistent with our study, suggesting that GPR180 may influence diabetes susceptibility or progression.

Despite these insights, several limitations should be acknowledged. The study relies on murine systems, which may limit extrapolation to human β-cell biology. Although the RIP-Cre model used for β-cell-specific deletion has been associated with off-target effects in some reports [33, 34], the consistent phenotype observed across all models supports a *bona fide* role for GPR180 in β-cells and reduces the likelihood of model-specific artifacts. While mitochondria emerged as a major downstream effector of GPR180, the precise mechanisms linking GPR180 to mitochondrial regulation remain incompletely defined. In addition, although MIN6 cells provide a well-established in vitro model to investigate β-cell biology [35, 36], their transformed nature may not fully recapitulate primary islet physiology.

Several important questions, therefore, remain. Future studies should define the molecular pathways downstream of GPR180 and determine how its activity is coordinated across metabolically relevant tissues. Notably, GPR180 expression has been reported in human pancreatic datasets, including GTEx and single-cell transcriptomic atlases, where it is detectable in β-cells at low to moderate levels [37, 38]. While this suggests potential conservation of GPR180 expression in human β-cells, functional validation in human islet models will be required to establish its relevance. Pharmacological modulation or rescue of GPR180 function *in vivo* could provide insight into its therapeutic utility.

Given its roles in both adipose tissue and β-cells, GPR180 may act as a broader coordinator of systemic energy metabolism. Future research should explore whether targeting GPR180 can simultaneously enhance insulin secretion and improve peripheral insulin sensitivity, potentially offering a dual-action strategy for metabolic disease.

## Data availability

The RNA sequencing data of murine islets, as well as MIN6 cells generated in this study, have been deposited in the NCBI Sequence Read Archive (SRA) under BioProject accession PRJNA1371626. Metabolomic data have been deposited in the MassIVE repository under accession number MSV000100260. Supporting data are included in this article as Supplementary data 1 and 2, Supplementary Figures 1 – 3 and Supplementary tables 1 and 2.

## Acknowledgement

We thank Manuel Klug (ETH Zurich) for his technical support with animal experiments and the SLA animal facility for assistance in mouse husbandry. We also acknowledge Ladislav Novota (Biomedical Research Center, Slovak Academy of Sciences) for expert assistance with the electron microscopy. The schematic overview of the GPR180 mechanism in the graphical abstract was created by modifying elements from Servier Medical Art (smart.servier.com) and BioRender.

## Authors’ relationship and activities

The authors declare that there are no relationships or activities that might bias, or be perceived to bias, their work.

## Contribution statement

L.B., C.W. and M.B. designed the study; L.B., T.D., O.K., D.F. and M.B. supervised the experiments; T.D., C.H. and L.B. conducted animal experiments; M.A., S.G., O.K., M.B. and L.B. performed *in vitro* functional studies; P.M. accomplished tissue histology; A.N. and M.N. realized ultrastructural analysis by electron microscopy; R.B., E.I., M.H. and D.F. conducted metabolomic study; D.S. and D.G. performed comprehensive searches for genetic variability of GPR180 by systematically querying publicly available databases; all authors contributed to data analysis and interpretation of the results; L.B. and C.W. obtained funding; L.B., M.A. and M.B. wrote the paper; all authors critically revised and approved the final version of the manuscript. L.B. is the guarantor of this work.

---

**Supplementary Figure 1.**
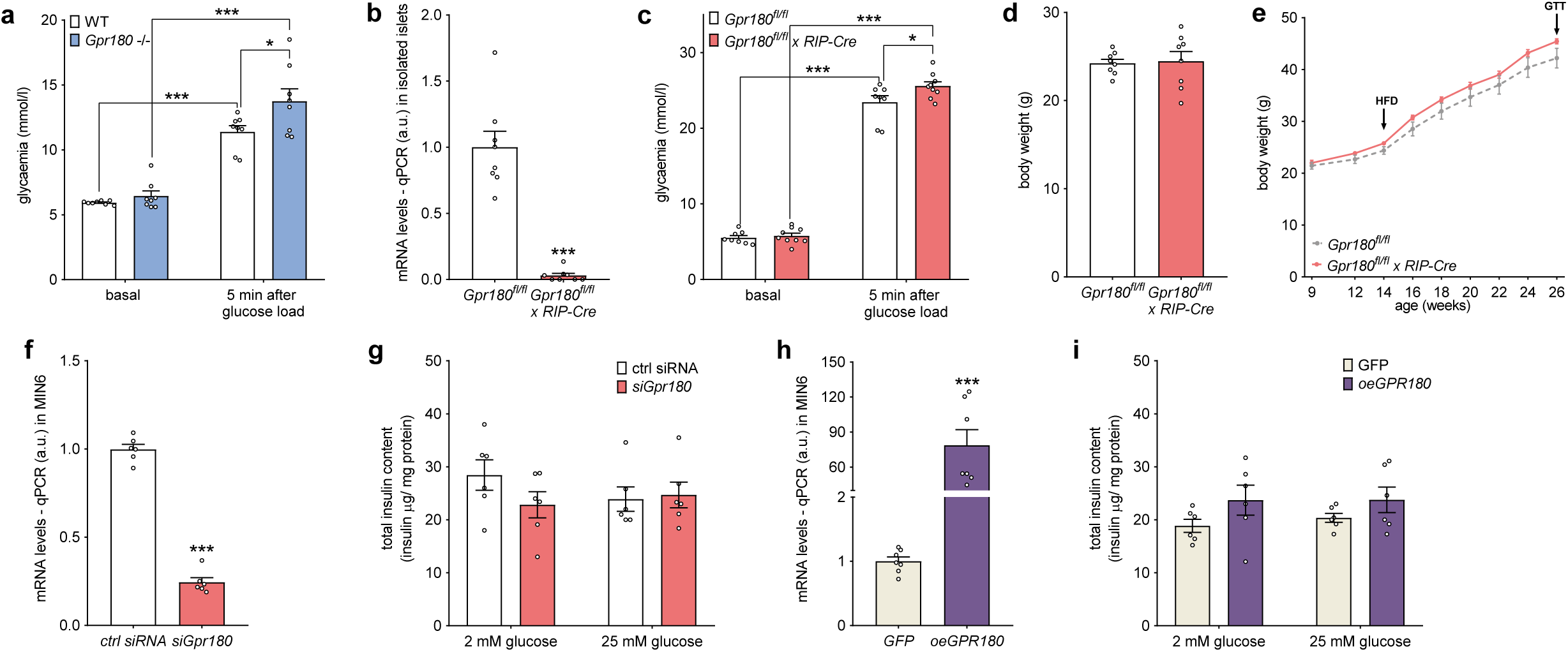
(Related to Figure 1): GPR180 deficiency impairs glucose-stimulated insulin secretion in a β-cell-autonomous manner. (a) Blood glucose level during the insulin secretion test in global *Gpr180* knockout mice and controls. (b) Knockout efficiency of *Gpr180* in isolated Langerhans islets from β-cell-specific *Gpr180* ablated mice. (c) Blood glucose level during insulin secretion test in β-cell-specific *Gpr180* knockout mice. Body weight in (d) chow- and (e) high-fat diet-fed mice with β-cell-specific *Gpr180* deletion (n = 8 for *Gpr180^fl/fl^*, n = 9 for *Gpr180^fl/fl^ x RIP-Cre*). (f) Knockdown efficiency of *Gpr180* in MIN6 cells following siRNA transfection. (g) Total insulin content in MIN6 cells after *Gpr180* silencing. (h) Validation of *Gpr180* overexpression in MIN6 cells following plasmid transfection. (i) Total insulin content in MIN6 cells after *GPR180* overexpression. Data are presented as mean ± SEM. Statistical analysis was performed by unpaired t-test (b, d, f, g), ordinary two-way ANOVA with subsequent Tukeýs post-hoc test (a, c), or two-way ANOVA with repeated measures followed by Sidak’s post hoc test (e). * p < 0.05; ** p < 0.01; *** p < 0.001. *Gpr180* G-protein coupled receptor 180, GTT glucose tolerance test, HFD high fat diet, RIP rat insulin promoter, WT wild type.

**Supplementary Figure 2.**
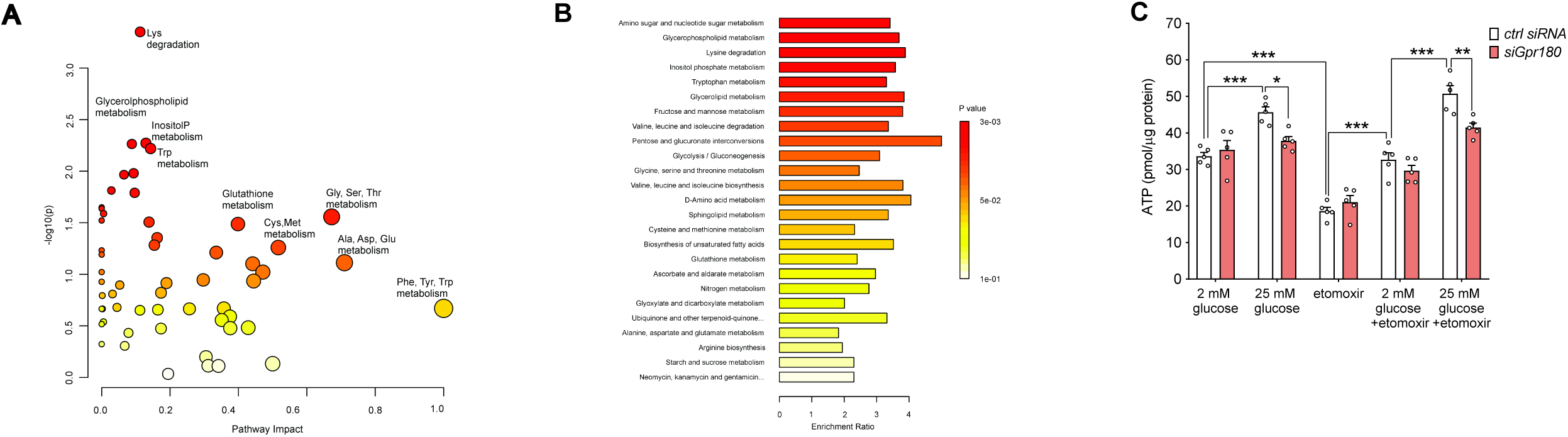
(Related to Figure 2): GPR180 affects insulin secretion by modulating mitochondrial glucose metabolism. (a) MetaboAnalyst pathway analysis of metabolic and organic acid datasets, showing significantly altered metabolic pathways based on differential metabolite abundance. (b) MetaboAnalyst enrichment analysis of metabolic and organic acid datasets, identifying metabolite classes and pathways over-represented among significantly changed metabolites. (c) ATP levels in MIN6 with siRNA-mediated *Gpr180* silencing in response to glucose and etomoxir (CPT1 inhibitor) or their combination. Data are presented as mean ± SEM. Statistical analysis was performed by ordinary two-way ANOVA with subsequent Tukeýs post-hoc test * p < 0.05; ** p < 0.01; *** p < 0.001. ATP adenosine triphosphate, *Gpr180* G-protein coupled receptor 180.

**Supplementary Figure 3.**
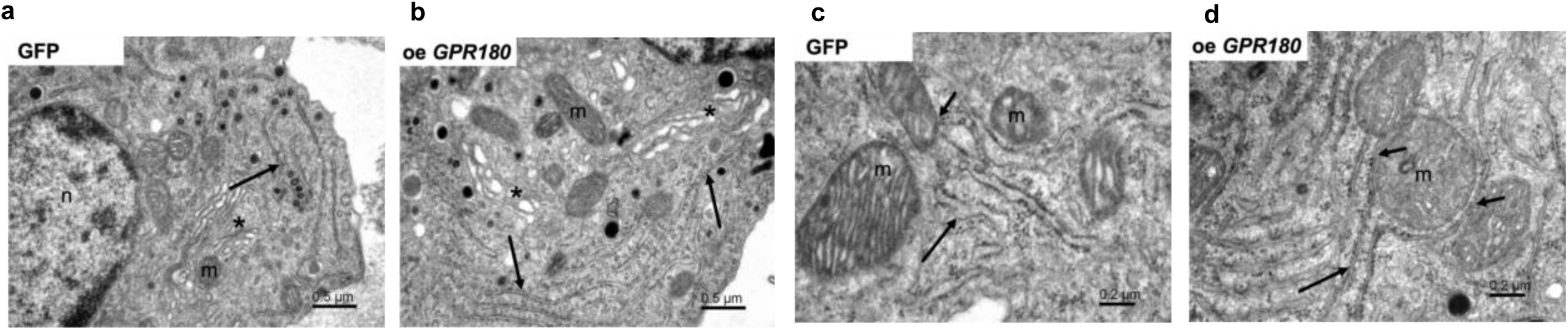
(Related to Figure 4): GPR180 alters cellular morphology and mitochondrial microenvironment. Representative transmission electron microscopy images of the cell overexpressing (a) GFP and (b) *GPR180*. Mitochondrial environment in (c) control GFP cells and (d) cells following *GPR180* overexpression. (a, b) scale bar 500 nm and (c, d) scale bar 200 nm. *Gpr180* G-protein coupled receptor 180, m - mitochondria, n - nucleus, asterisk - Golgi apparatus, long arrows - endoplasmic reticulum and short arrows - endoplasmic reticulum in proximity to the outer mitochondrial membrane

