## Supplementary Table 1 for "GPR180 deficiency impairs mitochondrial function and insulin secretion in pancreatic β-cells"

| **gene** | **Fwd primer sequence** | **Rev primer sequence** |
| --- | --- | --- |
| *Gpr180* | CGCAGTCTTCATCGTCATCA | CTGTGGTGACTGGTCTCTTCA |
| *GPR180* | GCATCGGCCACTTCGAGTTC | TGGGCTTGGAACAGGTAGAGT |
| *gLpl* | GGATGGACGGTAAGAGTGATTC | ATCCAAGGGTAGCAGACAGGT |
| *mt-Nd1* | CAGCCTGACCCATAGCCATAATAT | TGATTCTCCTTCTGTCAGGTCGAA |
| *Tbp* | GAAGCTGCGGTACAATTCCAG | CCCCTTGTACCCTTCACCAAT |

**Supplementary Table 1: List of primers used in qPCR.**
