## Supplementary Table 2 for "GPR180 deficiency impairs mitochondrial function and insulin secretion in pancreatic β-cells"

| **target** | **dilution** | **vendor** | **RRID** |
| --- | --- | --- | --- |
| AMPK, phosphorylated at Thr172 | 1 : 1000 | Cell Signaling | AB_331250 |
| AMPK, total | 1 : 1000 | Cell Signaling | AB_10622186 |
| gamma tubulin | 1 : 10 000 | Sigma–Aldrich | AB_477584 |
| HSP90 | 1 : 1000 | Cell Signaling | AB_2233307 |
| OXPHOS | 1 : 500 | Abcam | AB_2629281 |
| [IRDye 680RD Donkey anti-Mouse IgG](https://rrid.site/data/record/nif-0000-07730-1/RRID:AB_10953628/resolver?q=*&i=rrid:ab_10953628-1010534) | 1:10 000 | LI-COR | AB_10953628 |
| [IRDye 800CW Goat anti-Rabbit IgG](https://rrid.site/data/record/nif-0000-07730-1/RRID:AB_621843/resolver?q=*&i=rrid:ab_621843-180455) | 1:10 000 | LI-COR | AB_621843 |

**Supplementary Table 2: List of antibodies used in western blot.**
